# Long-term intrahost evolution of methicillin resistant *Staphylococcus aureus* among cystic fibrosis patients with respiratory carriage

**DOI:** 10.1101/599431

**Authors:** Taj Azarian, Jessica P. Ridgway, Zachary Yin, Michael Z. David

## Abstract

*Staphylococcus aureus* is the most commonly identified airway colonizer of cystic fibrosis (CF) patients, and infections with methicillin-resistant *S. aureus* (MRSA) are associated with poor outcomes. Yet, little is known about the intrahost evolution of *S. aureus* among CF patients. We investigated convergent evolution and adaptation of MRSA among four CF patients with long-term respiratory carriage. For each patient, we performed whole-genome sequencing on an average of 21 isolates (range: 19-23) carried for a mean of 1,403 days (range: 903-1,679), including 25 pairs of isolates collected on the same day. We then assessed intrahost diversity, population structure, evolutionary history, and signatures of adaptation in the context of patient age, antibiotic treatment, and co-colonizing microbes. We conducted tests of gene-wide diversifying selection and identified instances of switched intergenic regions (IGRs), which may be associated with gene expression. Phylogenetic analysis delineated distinct multilocus sequence type ST5 (n=3) and ST72 (n=1) clonal populations in addition to sporadic, non-clonal isolates, and uncovered a putative transmission event. Variation in antibiotic resistance was observed within clonal populations, even among isolates collected on the same day. Coalescent analysis revealed rates of molecular evolution ranging from 2.21 to 8.64 nucleotide polymorphisms per genome per year. Estimated lineage ages were consistent with acquisition of colonization in early childhood and subsequent persistence of multiple sub-populations. Selection analysis focused on 1,622 core genes shared by all four clonal populations (n=79). Eleven genes were variable in three subjects - most notable, ATP-dependent protease *clpX*, 2-oxoglutarate dehydrogenase *odhA, fmtC*, and transcription-repair coupling factor *mfd*. Only one gene, staphylococcal protein A (*spa*), was found to have evidence of gene-wide diversifying selection. We identified three instances of intrahost IGR switching events, two of which flanked genes related to quorum sensing. The ecology of microbes in the airways of CF patients is undeniably complex, posing challenges for management. We illustrate appreciable intrahost diversity as well as persistence of a dominant lineage. We also show that intrahost adaptation is a continual process, despite purifying selective pressure, and provide targets that should be investigated further for their function in CF adaptation.

**Contribution to the Field:** *Staphylococcus aureus* is the most commonly identified airway colonizer of cystic fibrosis (CF) patients, and infections with methicillin-resistant *S. aureus* (MRSA) are associated with poor outcomes. Yet, little is known about the intrahost evolution of *S. aureus* among CF patients. We investigated convergent evolution and adaptation of MRSA among four CF patients with long-term respiratory carriage. We found that each patient possessed a single predominant strain in addition to transient non-clonal isolates. Considerable genomic diversity was observed among intrahost populations including variation in antibiotic resistance, even among isolates collected on the same day. This finding has implications for treatment. Estimated lineage ages were consistent with acquisition of colonization in early childhood and subsequent persistence of multiple sub-populations. The scarcity of non-synonymous substitutions suggests chronic *S. aureus* carriage in CF patients provides a sufficient opportunity for purifying selection to act. Eleven genes were found to be variable in three patients and present possible candidates for loci experiencing gene-wide diversifying selection. Overall, we show that intrahost adaptation is a continual process, despite purifying selective pressure, and provide targets that should be investigated further for their function in CF adaptation.

## Introduction

Cystic fibrosis (CF) is an autosomal recessive, debilitating disease that is caused by mutations in the gene encoding the CF transmembrane conductance regulator protein (CFTR), which occurs at an incidence of 1/2,500 live births. CF manifests as a clinical syndrome characterized by insufficient mucociliary clearance with chronic pulmonary infections that ultimately leads to a progressive decline in lung function and shortened life span of these patients. A key component of CF patient treatment is management of bacterial respiratory pathogens, including *Staphylococcus aureus*, which may colonize the CF lung for years (Branger et al., 1996). This often requires recurrent and prolonged periods of antimicrobial use (Schwerdt et al., 2018). Together with the frequent occurrence of polymicrobial carriage and infection, host immunity, and the challenges of nutrient acquisition, the evolutionary forces shaping intrahost microbial populations colonizing the CF lung are complex (McAdam et al., 2011). Identifying important genotypes of pathogenic bacterial species that are associated with adaptation to persistence and deciphering which genotypes represent bystanders, protective commensals, or pathogens, are central to long-term clinical management.

Among persistent colonizers of the CF lung, *Staphylococcus aureus* may cause recurrent infections (Harrison, 2007; Kahl, 2010). However, in comparison to *Pseudomonas aeruginosa* (Feliziani et al., 2014; Marvig et al., 2015) or even *Burkholderia* spp.(Lieberman et al., 2011a, 2014; Price et al., 2013; Silva et al., 2016), less is known about *S. aureus* intrahost evolution in the CF lung (Ankrum and Hall, 2017; McAdam et al., 2011; Rolain et al., 2009; Schwerdt et al., 2018). The various species inhabiting the CF lung share several evolutionary strategies relating to persistent microbial carriage. Persistent colonization usually involves a single clonal population with the sporadic appearance of non-clonal isolates. After initial colonization there is a rapid increase in population size as the microbe adapts to the lung environment (Lieberman et al., 2011b). This adaptation usually involves resistance to antimicrobials through target site modification or gain of mobile genetic elements carrying antibiotic resistance associated genes as well as substitutions in metabolic loci and the formation of small colony variants or hypermutators (Feliziani et al., 2014). Local adaptation may also occur in various compartments of the respiratory system, resulting in intrahost sub-population structure (Lieberman et al., 2014). Drift, population structure, and relaxed purifying selection often lead to evolutionary rates that are higher than those observed in studies of interhost transmission (Didelot et al., 2016). Yet, despite these shared evolutionary strategies of bacteria in the CF lung, there remains tremendous variability in observations across pathogens and studies.

Here we investigate the intrahost diversity, host adaptation, and convergent evolution of methicillin-resistant *S. aureus* (MRSA) among four CF patients with long-term respiratory carriage. For each patient, we performed whole-genome sequencing on an average of 21 isolates (range: 19-23) carried for a mean of 1,403 days (range: 903-1,679). We sought to identify trends in population dynamics within and between patients, specifically assessing diversifying selection in the same genes across clonal intrahost populations, which would provide the strongest evidence for convergent evolution. We further consider variation in intergenic regions (IGR) and the impact of within-host IGR switching events associated with regulatory modification.

## Materials and Methods

### Bacterial strains, genome sequencing, and assembly

We investigated *S. aureus* isolates collected from four patients who were seen at the CF clinic at the University of Chicago from 2003 to 2009 (see Table S1 in the supplemental material). Clinical, laboratory, and demographic data were abstracted from the medical records. This included the antibiogram for each isolate as determined by Vitek, age, recent hospitalizations, presence of indwelling devices, recent surgery, antibiotic treatment, identification of other respiratory organisms, and lung function tests, when available. Informed consent was obtained from participants 18 years of age or older; for participants between 7 and 17 years of age, consent was obtained from the parent or guardian as well as assent from the child. The study was approved by the Institutional Review Board of the Biological Sciences Division of the University of Chicago.

Genomic DNA (gDNA) was extracted from clinical isolates using the Qiagen blood and tissue kit (Qiagen). WGS libraries were constructed using the Nextera XT kit, which were subsequently sequenced on an Illumina HiSeq to produce 2×150 paired end reads. *De novo* assembly was performed with Spades v3.11.1 followed by scaffolding with SSPACE v3.0. Assembly refinement (gap filling and error correction) was carried out using three iterations of Pilon v1.12. Final assemblies were filtered to remove contigs shorter than 250 bp and censor sites with ≤ 5 read depth, base quality ≤ 20, and mapping quality ≤ 20. Genome annotation was then performed using Prokka v1.13. Multilocus sequence types (MLST) were assigned using the Center for Genomic Epidemiology web server (https://cge.cbs.dtu.dk/), which also identified the closest matching reference strain in NCBI (Zankari et al., 2013). Antibiotic resistance associated genes were identified using ARIBA v2.12.1 and the CARD database (Hunt et al., 2017). Pan-genome analysis was performed on the entire sample and multiple references using Roary v3.12.0 (Page et al., 2015). A single-nucleotide polymorphism (SNP) alignment was extracted from the core-genome alignment using snp-sites v2.4.0 and a maximum likelihood phylogeny was inferred with RAxML v8.2.11 using the ASC_GTRGAMMA substitution model with 100 bootstrap replicates (Stamatakis, 2014).

### Inter- and intrahost population structure

The ML phylogeny was used to assess inter- and intrahost population structure to determine the clonality of each patient’s *S. aureus* population and identify possible epidemiologic linkages (i.e., transmission events). Mean pairwise SNP distances were calculated for each intrahost population. Isolates with divergent MLST types, as compared to the major frequency ST, were excluded, as well as isolates belonging to the major frequency ST with mean pairwise SNP distances greater that three standard deviations of the mean. These later isolates were indicative of independent acquisitions of *S. aureus* unrelated to the dominant carried population. For clonal inter- and intra-host *S. aureus* populations, pan-genome analysis was repeated, core-genome alignments were obtained, and ML phylogenies inferred. SNPs in the core genome were then used to assess intrahost population structure with RhierBAPS (Cheng et al., 2013; Tonkin-Hill et al., 2018). Last, Piggy v1.4 was used to assess intergenic regions (IGR) and identify “switched” IGRs upstream of genes, which could impact gene expression (Thorpe et al., 2018).

### Intrahost diversity and coalescent analysis

Intrahost variation was explored through assessment of allele frequency and SNP diversity compared to collection time point. For each intrahost population, the temporal signal, indicative of a measurably evolving population, was investigated by regressing the root-to-tip genetic distance against sampling times. To maintain confidentiality, all dates have been scaled as years from collection of the first isolate in the study sample (i.e., Year 0). We used BacDating v1.0 to estimate the intrahost evolutionary rates and date the most recent common ancestor (MRCA) (Didelot et al., 2018). MCMC chain lengths were run for 100-800 million until effective sample sizes (ESS) over 100 were observed for *mu, sigma*, and *alpha* parameters, as suggested by the authors. MRCAs were then compared to clinical and demographic characteristics such as age and time of follow-up.

### Selection analysis

We investigated signatures of selection among inter- and intra-host populations, focusing specifically on core genes present among all isolates in the clonal intrahost sample. First, for each core gene, we calculated mean pairwise SNP diversity and Watterson’s Θ at 0-fold and 4-fold degenerate sites, i.e., those nucleotide sites which all changes are synonymous and non-synonymous, respectively. We then assessed gene-by-gene diversity across all intrahost populations and focused subsequent analysis on genes that demonstrated nucleotide variation in at least two patients’ intrahost populations. For those genes, we used the tool BUSTED (Branch-Site Unrestricted Statistical Test for Episodic Diversification) implemented in HyPhy v2.3.14 to test for gene-wide diversifying selection on internal branches of the genealogy (Murrell et al., 2015; Pond et al., 2005). BUSTED analysis was carried out on the entire interhost sample as well as each clonal intrahost population. This allowed for the identification of genes potentially experiencing diversifying selection during the duration of intrahost carriage as well as at a population level. Functional annotations for genes experiencing selection were assessed using Kegg Orthologies (www.kegg.jp).

## Results

Patient age at the start of follow-up ranged from 4 to 26 years of age with a duration of follow-up ranging from 903 to 1,759 days (mean=1,403 [3.8 years]) (Table 1). For patients one through four, we analyzed 21, 23, 22, and 19 *S. aureus* isolates, respectively, of which 18, 22, 21, and 18 were identified as clonal based on MLST typing, SNP divergence, and interhost phylogeny (Figure 1). Duration of time between isolates ranged from 0 days (collected on the same day, n=24 pairs) to 547 days, with the majority of isolates collected during routine quarterly visits (average time between isolate pairs = 69.28, standard deviation = 87.0, median = 49, Supplementary Figure 1). Three patients (1, 2, and 4) were colonized with a dominant ST5 population while Patient 3 was colonized with ST72. Non-clonal isolates often belonged to ST8, the most prevalent community-associated MRSA lineage in the US (Otter and French, 2012). In each instance, the ST8 isolate was only observed at a single time point. In general, there was little evidence of recent transmission or intermixing of clonal populations. However, in one instance of a putative epidemiological linkage, two ST5 isolates belonging to Patients 1 and 2 appeared divergent from their clonal populations yet identical to each other (0 core genome SNP differences). Closer inspection of the collection dates revealed that these isolates were collected 37 days apart. No other instances of suspected transmission events were observed.

**Table 1.** Patient demographic information and *S. aureus* molecular characteristics.

| Patient | Age at start | Duration of follow-up (days/years) | Number of isolates | Clonal isolates | MLST of primary clone | Other MLSTs |
| --- | --- | --- | --- | --- | --- | --- |
| 1 | 9 | 1,679 / 4.6 | 21 | 18 | ST5 | ST8, ST2232 |
| 2 | 18 | 1,271 / 3.5 | 23 | 22 | ST5 | ST8 |
| 3 | 4 | 1,759 / 4.8 | 22 | 21 | ST72 | ST5 |
| 4 | 26 | 903 / 2.5 | 19 | 18 | ST5 | ST8 |

**Figure 1.**
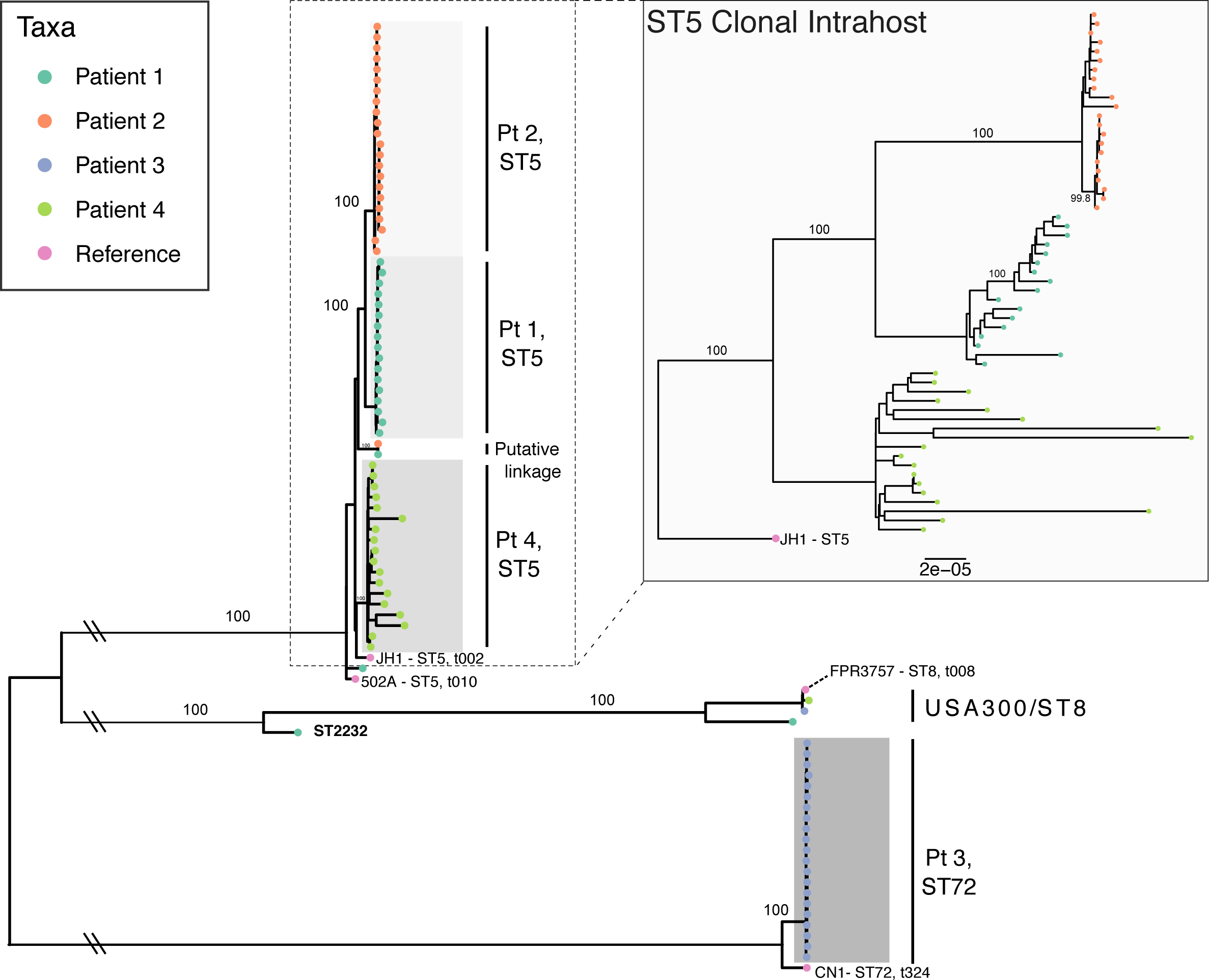
Maximum likelihood phylogeny of all intrahost isolates from four patients with *S. aureus* reference genomes. The main phylogeny was inferred using RAxML from a core genome alignment of all intrahost isolates with four *S. aureus* reference genomes (JH1, 502A, FPR3757, and CN1), which are annotated with multilocus sequence type (ST) and *spa* type. Tip points are colored by patient and bootstrap values for major clades are annotated. Clades are shaded to identify clonal intrahost populations, which are labeled to the right. The inlayed phylogeny shows ST5 clonal intrahost isolates only and was inferred from a core gene alignment of 2,164 COGs (1,966,902 bp) rooted using the JH1 ST5 reference.

We assessed nucleotide diversity in the core genome and IGRs of each clonal intrahost population. Intrahost nucleotide diversity varied significantly among subjects, regardless of duration of follow-up and core genome size (Table 2). In particular, Patient 3 with the ST72 population had the lowest diversity in both core and IGR despite the longest duration of follow-up (4.8 years), while Patient 4, who had the shortest duration of follow-up (2.5 years), had the highest core and IGR nucleotide diversity. Variation was also observed in the distribution of SNP frequencies (Supplemental Figure 2). At opposite extremes, Patient 2 had a higher density of intermediate frequency SNPs, while Patient 4 had a higher density of singletons. In general, pairwise SNP distances generally increased with time separating the collection date of isolates (Supplemental Figure 1). This is consistent with our observation when we assessed root-to-tip phylogenetic distance to determine temporal signal (see coalescent analysis below). We found a significant range in intrahost diversity between 25 pairs of isolates collected on the same day (Supplemental Figure 3). The amount of diversity among isolates collected on the same day could in part be explained by the observation that these pairs were often collected from patients later in their follow-up (Mean=904.2 days), reflecting larger intrahost population sizes during persistent colonization or the existence of hypermutator strains that evolved later in carriage. Overall, there was a significant range in intrahost diversity among same-day isolates from each patient. Last, assessment of clonal intrahost population structure identified three to four ‘sequence clusters’ within each population (see phylogenies and coalescent analysis below); however, due to the low diversity of each intrahost population, we are cautious about the interpretation here.

**Table 2.** Intrahost core-genome and intergenic (IGR) diversity of clonal *Staphylococcus aureus* isolates. Diversity estimates represent mean pairwise single-nucleotide polymorphism (SNP) differences with standard errors (SE).

| Patient | Core Genes | Core genome size (bp) | Variable Core Genes | Core SNP Diversity (mean, SE) | Variable IGR | IGR SNP Diversity (mean, SE) |
| --- | --- | --- | --- | --- | --- | --- |
| 1 | 2263 | 2,008,627 | 73 | 40.2 (1.84) | 13 | 4.10 (0.18) |
| 2 | 2469 | 2,218,136 | 57 | 44.6 (2.30) | 15 | 4.17 (0.18) |
| 3 | 2405 | 2,196,755 | 16 | 16.4 (0.83) | 4 | 2.5 (0.11) |
| 4 | 2125 | 1,938,257 | 125 | 181.2 (9.04) | 78 | 19.99 (1.03) |

**Figure 2.**
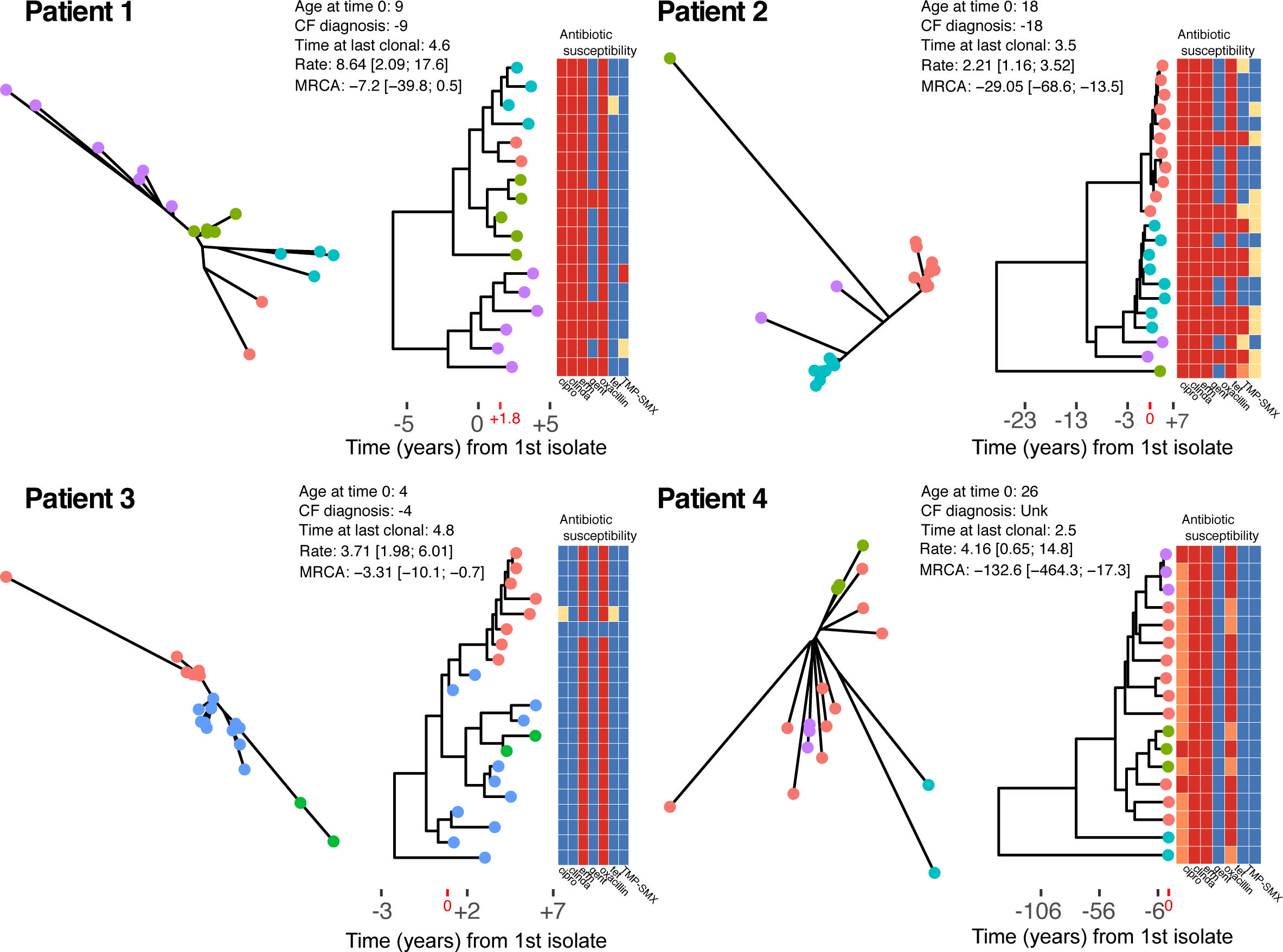
Un-rooted maximum likelihood phylogenies (left), and time-scaled Bayesian coalescent phylogenies with heatmap of phenotypic antibiotic susceptibilities (right) for each patient. Tip labels are colored by sequence cluster as identified by rhierBAPS and illustrate inferred intrahost sub-population structure. Time is scaled as years from the date of the first isolate in our study sample (Year 0), and times are provided for the CF diagnosis, age at time 0, and time at the date of the last clonal isolate (i.e., years of follow-up). Evolutionary rates scaled as SNPs/core-genome/year and estimates of time to the most recent common ancestor (years) are annotated on the phylogeny. The heatmap shows the phenotypic susceptibility results (resistant - red, intermediate - orange, susceptible - blue, and not tested - yellow) for ciprofloxacin (cipro), clindamycin (clinda), erythromycin (erm), gentamicin (gent), oxacillin, tetracycline (tet), and trimethoprim-sulfamethoxazole (TMP-SMX).

**Figure 3.**
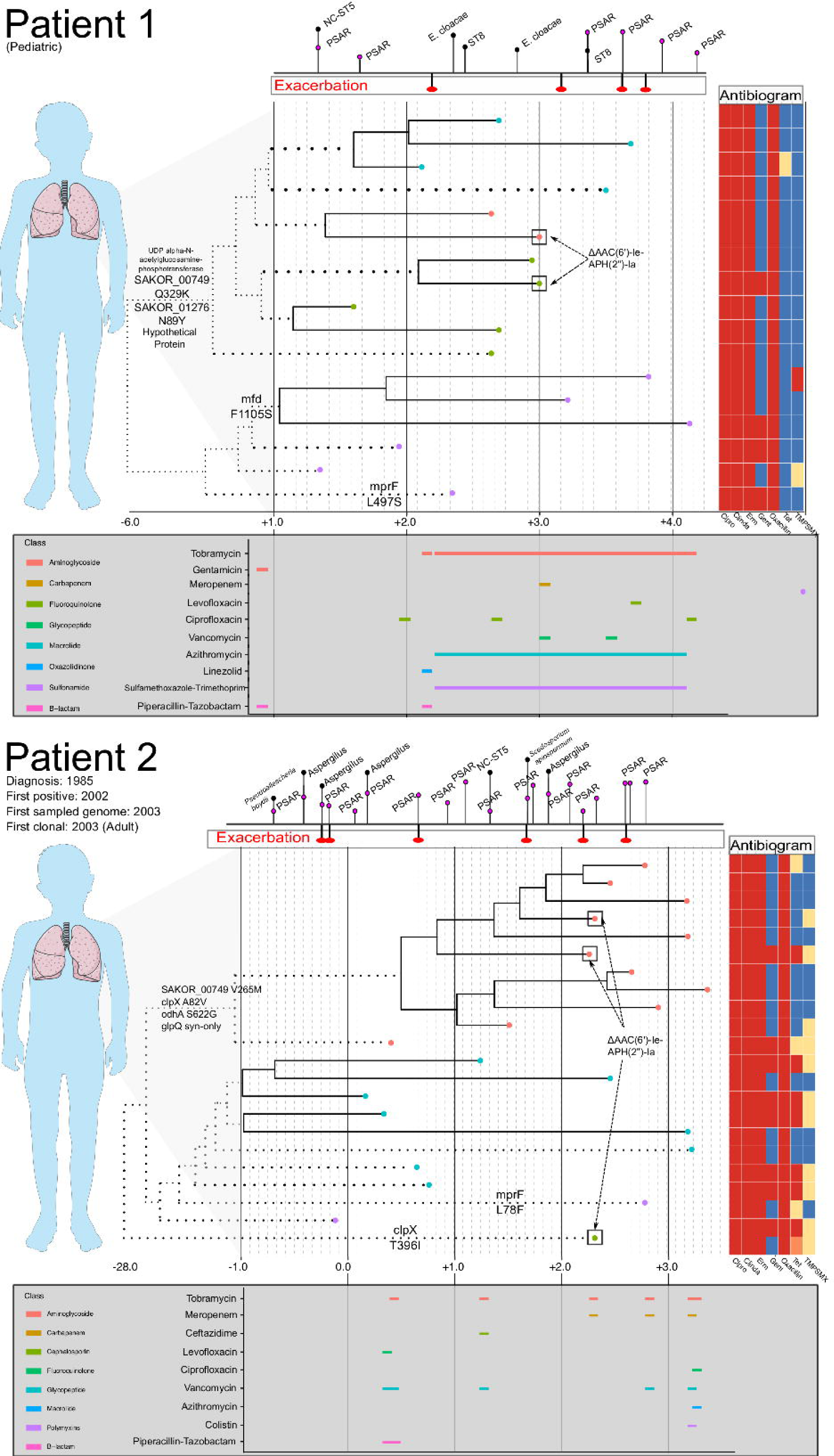
Intrahost genomic epidemiology of Patients 1 and 2. For Patients 1 and 2, a time-scaled phylogeny (as shown in Figure 2) is shown, annotated with phenotypic antibiogram, antibiotic treatment, and notable genomic events. Time is scaled as years from the date of the first isolate in our study sample (Year 0). The phylogeny has been compressed to focus on events occurring during the study period and branches that have been scaled are dotted. Tip labels are colored based on intrahost sub-population structure. Above each time-series is a timeline dating CF exacerbations and identification of other respiratory organisms including *Pseudomonas aeruginosa* (PSAR) and non-clonal (NC) *Staphylococcus aureus*. Below each time series is a timeline showing antibiotic usage and drug class (legend to the left of the timeline) during the study period. For Patient 1, chronic azithromycin and trimethoprim/sulfamethoxazole was prescribed three times a week. Inhaled tobramycin was prescribed every other month (every 12 hours, 28 days on, 28 days off). Instances where antibiograms varied among isolates collected on the same day or within 14 days are annotated on the tree. Last, notable substitutions among loci listed in Table 3 are annotated on the branches of the phylogeny, specifically focusing on substitutions that occurred on basal internal branches that bifurcate dominant clades of the intrahost population. Patients 3 and 4 were not included because of lack of completeness of their treatment and clinical history.

### Intrahost coalescent analysis

We sought to estimate intrahost evolutionary rates and date the MRCA for each patient’s clonal *S. aureus* population. First, temporal signal was evaluated by assessing the root-to-tip collection using maximum likelihood phylogenies and the day of isolation. For all clonal intrahost populations, root-to-tip correlations were significant; however, correlation coeffecients were relatively low for Patients 2 and 4 (Supplemental Figure 4). This was likely the result of outliers in the root-to-tip analysis as well as the existence of intrahost sub-population structure. Evolutionary rates ranged from 2.21 [95% highest posterior densities (HPD): 1.16, 3.52] to 8.64 [2.09, 17.6] SNPs per year in the core-genome (Figure 2). We further investigated the gene sequence of DNA mismatch repair protein *mutS* and *mutL*, which have previously been associated with hypermutator phenotypes (Prunier and Leclercq, 2005). We identified non-synonymous mutations in both genes among patient 4 isolates, suggesting a possible role in high intrahost diversity.

**Table 3.**
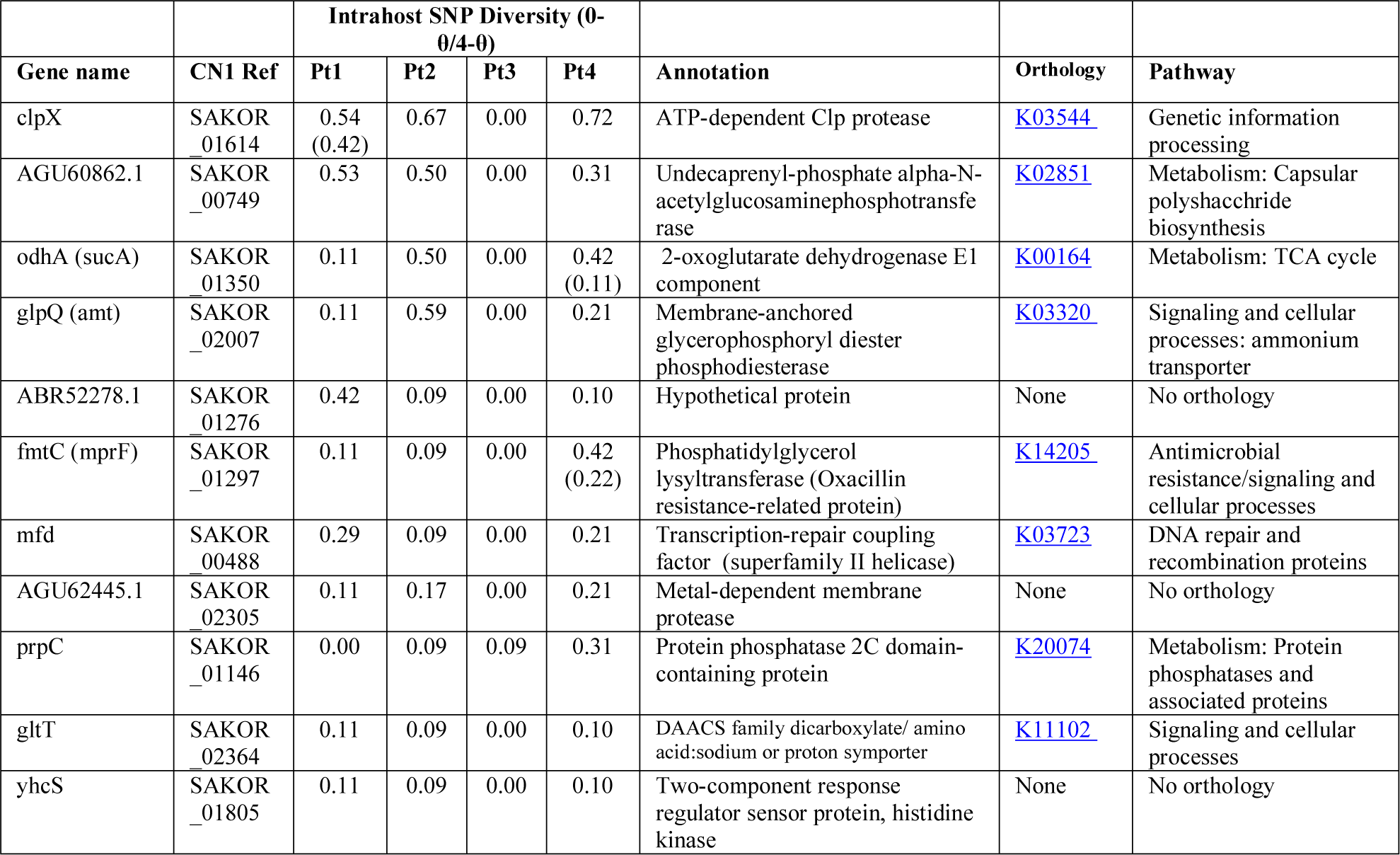
Core genome clusters of orthologous groups (COGs) that contained nucleotide diversity among three clonal intrahost populations. Gene names are provided where available; alternatively, the locus name in the CN1 reference is given (NCBI: CP003979.1) as well as the locus tag (CN1 Ref). Intrahost diversity represents the mean pairwise single-nucleotide polymorphism (SNP) difference. Where calculable, the 0-fold vs. 4-fold θ values are included with the diversity estimates. The annotation, orthology, and pathway from Kegg (www.kegg.jp) are included for each gene.

In general, younger individuals (Patients 1 and 3) had MRCAs that were consistent with primary colonization by a clonal population at a young age. In the case of Patient 3, *S. aureus* was likely acquired during infancy. Older individuals (Patients 2 and 4) had MRCAs that predated their biological age. Specifically, Patient 1 was diagnosed in infancy and had the first positive *S. aureus* isolate five years latter. The earliest isolate in our sample (Year 0) was collected four years after the first *S. aureus* isolate, and the first clonal isolate was observed nearly two years later (+1.8) (Figure 2). Coalescent analysis dates the MRCA of the clonal population near the time of diagnosis. Patient 2 was 18 years of age when the first isolate in our sample was obtained (Year 0), with the first positive *S. aureus* respiratory culture documented one-year prior. The MRCA for the clonal population was estimated prior to their date of CF diagnosis. Patient 3 was diagnosed with CF in infancy and had the first positive *S. aureus* respiratory culture two years later, three years prior to the first isolates in our sample (Year 0), which was collected at 4 years of age. Coalescent analysis dated the MRCA for the clonal population as −3.3 [95% HPD: −10.1,-0.7]. Patient 4, an adult CF patient, has an unknown diagnosis date and had the first positive *S. aureus* respiratory culture at the age of 26, with the first isolate in our sample collected that year. Our follow-up for Patient 4 began with two isolates that were collected on the same day, which were separated by 131 SNPs, demonstrating appreciable intrahost diversity within a clearly defined clonal population. Consistent with the high intrahost diversity, coalescent analysis dated the MRCA at over a hundred years, with two basal isolates driving the estimate (Figure 2D). Overall, despite evolutionary rates with over-lapping HPDs (i.e., not significantly different), intrahost diversity and corresponding MRCA estimates varied among patients and was associated with age. This may reflect differences in mutation rates or the amount of selective pressure on pathogen populations among patients and the time those forces were acting on the populations. To investigate this further, we searched for signals of diversifying selection within and among intrahost populations.

### Antibiotic resistance

Within clonal intrahost populations, we found sporadic variation in phenotypic antibiotic susceptibility. ST5 populations in patients were generally resistant to fluoroquinolones, lincosamides, macrolides, and penicillin, while Patient 4’s ST72 population was only resistant to macrolides and penicillin. Variation in aminoglycoside (i.e., gentamicin) and tetracycline resistance was the most common, occurring in Patients 1 and 2 and not isolated to specific clades of the population (Figure 2). We found that gentamicin resistance was conferred by AAC(6′)-Ie-APH(2″)-Ia aminoglycoside acetyltransferase (ARO:3002597), which was harbored on a 3kb transposon. In at least one instance in an isolate from Patient 2 (CF_Pt2_3_166), we found that the transposon was inserted in a 2.2kb plasmid. However, it is difficult to conclude from the genome assemblies if this was consistent across all isolates possessing the transposon. During their clinical management, Patients 1 and 2 routinely received the aminoglycoside tobramycin, which could have driven resistance although in Patient 2, the clade that predominantly harbored gentamicin resistance (lower clade with teal-colored tip points in Figure 2) was later replaced by a predominantly susceptible clade. Unfortunately, the absence of detailed antibiotic treatment data for Patients 3 and 4 prevents a more in-depth analysis of the impact of antibiotic use on intrahost population structure.

### Selection analysis

Selection analysis primarily focused on 1,622 core genes (1,484,432 bp) shared by all four clonal intrahost populations as well as two lineage specific references, JH1 for ST5 and CN1 for ST72, which allowed for annotation of functional pathways. First, we assessed nucleotide diversity (Watterson’s Θ) at 0-fold (strictly non-synonymous) and 4-fold (synonymous) degenerate sites in each clonal intrahost population, finding low 0-fold/4-fold values for the few genes for which this statistic could be calculated. Only three of 71 estimates were above one (Supplemental Figure 4).

Supplemental Table 2 lists the clusters of orthologous groups (COGs) variable in two or more patients with annotations and population genetic statistics, i.e., those genes in which mutations arose during intrahost colonization. Within the core genome of the clonal intrahost sample (n=79), eleven COGs were variable in three subjects (Table 3) and 119 were variable in at least two isolates. There were no COGs that were variable among all four patients, and most often, variable COGs were observed among ST5 isolates belonging to Patients 1, 2, and 4. Among these genes were ATP-dependent protease *clpX*, 2-oxoglutarate dehydrogenase E1 *odhA, fmtC*, and transcription-repair coupling factor *mfd*.

Selection analysis performed on all (ST5 and ST72) clonal isolates using BUSTED identified seven COGs that were significantly associated (p-value < 0.05) with gene-wide episodic diversifying selection among internal branches in the phylogeny (Supplemental Table 2). BUSTED analysis was then run using alignments from each patient’s clonal population as well as on only the patients with ST5 clonal populations. This represented the strictest test for diversifying selection, requiring substitutions to have occurred intrahost and selection to act upon those amino acid changes. Only one gene, staphylococcal protein A (*spa*), was found to have evidence of gene-wide diversifying selection in a patient (Patient 4, p-value = 0.025) and the ST5 sub-sample (Patients 1/2/4, p-value = 0.0002).

Last, we examined intrahost IGR diversity and occurrences of intrahost IGR switching events, which have recently been shown to correlate with changes in gene expression (Thorpe et al., 2018). We identified 611 core IGR (303,529 bp) found among all four clonal intrahost populations and lineage specific references. Of those, six were diverged in two or more intrahost samples. Three of these instances were the result of divergence between ST5 and ST72; however, in one instance we observed the same mutation arise intrahost. This occurred in the IGR flanking the coding sequences for phenol soluble modulin β (PSMb) (SaurJH1_1257 and SaurJH1_1258) in Patients 1 and 4, who shared the C->T mutation at position 40 of the IGR. There were three instances of intrahost IGR switching events, one each occurring in three patients (Table 4). While the phenotypic impact and significance of these switches remains to be determined, two switches flanked genes related to quorum sensing.

**Table 4.** Intrahost intergenic (IGR) switching events. Gene annotations are provided using the closest reference to each patient’s clonal reference isolate (CN1, CP003979.1, for ST72 isolates and JH1, CP000736.1, for ST5 isolates).

| Patient | Isolates Pairs | Days separating sampled isolates | IGR Length (bp) | Gene orientation | Flanking gene 1 | Annotation | Flanking gene 2 | Annotation | Pathways |
| --- | --- | --- | --- | --- | --- | --- | --- | --- | --- |
| 1 | CF_Pt1_20_1574 - CF_Pt1_10_1142 | 432 | 819 | Double promoter | ABR53315 (SaurJH1_2491) | Cobalt-zinc-cadmium resistance protein, czcD | ABR53316 (SaurJH1_2492) | Immunoglobulin G-binding protein Sbi | Infection |
| 3 | CF_Pt3_18_1503 - CF_Pt3_22_1759 | 256 | 364 | Co-oriented forward | AGU61240 (SAKOR_01100) | Phenol-soluble modulins beta | AGU61241 (SAKOR_01101) | HAD-superfamily hydrolase, subfamily IA | Quorum sensing |
| 4 | CF_Pt4_9_688 - CF_Pt4_11_773 | 85 | 712 | Double terminator | ABR52993 (SaurJH1_2164) | Metal dependent hydrolase | ABR52994 (SaurJH1_2165) | YidC/Oxa1 family membrane protein insertase | Quorum sensing, Protein export, Secretion system |

## Discussion

*S. aureus* overtook *P. aeruginosa* as the most prevalent respiratory organism among American CF patients in 2003 (Cystic Fibrosis Foundation Patient Registry, 2017). Since then, the prevalence of *S. aureus* and MRSA has continued to increase, with more than half of all CF patients having at least one positive *S. aureus* culture in 2017. This trend is concerning as MRSA been independently associated with decreased lung function and increased mortality among CF patients (Dasenbrook et al., 2008, 2010). *S. aureus* is also one of the earliest colonizers of the CF airway, and acquired strains can persist for years, often dominated by a single intrahost lineage (Al-Zubeidi et al., 2014; Kahl et al., 2003; Schwerdt et al., 2018). However, trajectories of *S. aureus* carriage vary by patient (Kahl et al., 2013; Schwerdt et al., 2018), reflecting the complexities of patient-specific CF management and complicating our understanding of pathogen adaptation. Despite its high prevalence, arguably far less is known about *S. aureus* genomic epidemiology and intrahost evolution than other CF pathogens such a *P. aeruginosa* or *Burkholderia* spp. Here, we studied the intrahost evolution of four patients with persistent MRSA carriage to identify trends in pathogen adaptation. We illustrate persistence of a dominant lineage as well as appreciable intrahost genomic diversity, including variation in antibiotic resistance, even among isolates collected on the same day. We also show that intrahost adaptation is a continual process, despite purifying selective pressure, and provide targets that should be investigated further for their function in CF adaptation.

We found that each patient studied here possessed a persistent clonal *S. aureus* population characterized by considerable genomic and phenotypic diversity as well as intrahost population structure. Within-host nucleotide diversity varied considerably among patients while intrahost evolutionary rates were consistent, ranging from 1.91E-06 to 9.96E-07 SNPs per site per year. These rate estimates fall within the range of 10^-7^ to 10^-5^ detailed in previous intrahost studies, considerably higher than interhost rates calculated over longer timescales (Biek et al., 2015; Didelot et al., 2016). We find that Patients 2 and 4, who were older at the start of the study than Patients 1 and 3, had significantly higher overall diversity estimates. Snapshot estimates of intrahost diversity gleaned from isolates collected on the same day also showed higher values for Patients 2 and 4. As a result, the MRCAs for their populations are estimated to be considerably older, and while these estimates were driven in both instances by phylogenetically basal outliers, they suggest that Patients 2 and 4 have likely been persistently colonized since early childhood or even infancy. Indeed, the MRCA estimates from Patients 1 and 3 are consistent with early colonization, matching epidemiological data from the general CF population that identifies *S. aureus* as one of the earliest colonizers (Cystic Fibrosis Foundation Patient Registry, 2017).

Intrahost population dynamics of *S. aureus* are likely to differ by patient due to a number of clinical and epidemiological factors that exert varied selective pressure. Here, further consideration of Patient 4’s clonal population in context of other ST5 isolates (Figure 1) suggests differences in the underlying population dynamics. Phylogenetic branch lengths are longer, consistent with higher diversity, and the unrooted ML phylogeny is more star-like (Figure 2). This contrasts all other patients, which have well-defined intrahost population structure - each with at least two extant lineages. While Patient 4 possessed mutations in genes previously associated with hypermutator phenotypes, it is more likely that a longer carriage duration and higher intrahost population size are responsible for the observed diversity since the estimated evolutionary rate was not significantly different. It is unclear whether Patient 4’s clonal population represents a single dominant intrahost lineage; however, the population structure within other patients reinforces previous findings that mutations rarely fix in intrahost pathogen populations. This suggests that intrahost clonal interference – whereby lineages are unable to reach fixation due to competition between multiple beneficial genotypes – is a common theme in intrahost population dynamics (Fogle et al., 2008; Gerrish and Lenski, 1998; Lieberman et al., 2014). Indeed, the distribution of SNP densities largely shows a high density of intermediate mutations that correspond to the population structure observed within each patient’s clonal population. Continued surveillance of intrahost populations in cases of chronic carriage may reveal whether these sub-populations persist or are removed in a bottleneck event.

In addition to each patient’s dominant clonal population, transient *S. aureus* strains were also identified during their course of follow-up. This finding was consistent with our previous study on intrahost evolution of *S. aureus* ST8 among individuals with recurrent skin and soft tissue infections, which found that non-ST8 strains often followed antibiotic treatment and preceded strain replacement (Azarian et al., 2016). Here, we observed that invading strains, which were often ST8 – the most common community-associated MRSA strain in the US – did not supplant the resident population. The same was also true for the two non-clonal ST5 isolates collected from Patients 1 and 2, which appeared to be epidemiologically linked. These strains were not phenotypically more susceptible to antibiotics, as may be expected, suggesting other dynamics at work. The resident population may be better adapted to persistence – for example, have attenuated virulence compared to invading ST8 stains – or other competitive advantage as has recently been described in asymmetric owner-intruder competitive strategies among pneumococci (Lees et al., 2018). In the latter, quorum-sensing mediated fratricide is employed by resident bacteria to exude ‘ownership’ of a host, resisting challenge by intruders. In staphylococci, the quorum-sensing accessory gene regulator (agr) system, which plays a role in biofilm formation and bacterial persistence, may fill a similar role and warrants further investigation (Painter et al., 2014). Overall, frequent healthcare contact increases exposure of CF patients to pathogens, and understanding why these sporadic strains fail to persist would assist in ascribing risk as well as further our knowledge or microbial interactions.

Persistence of *S. aureus* among CF patients is well recognized, and long-term carriage is often associated with specific phenotypes including small-colony variants, antibiotic resistance, and biofilm-formation (Branger et al., 1996; Dasenbrook et al., 2008; Kahl et al., 1998, 2003). Persistence generally involves the ability to grow in nutrient poor conditions, resist chronic antibiotic use, evade host defenses, tolerate stress, and attenuate virulence (Richards et al., 2015). Here were characterize persistence and assess the selective pressures shaping genomic variation associated with these traits, specifically investigating convergent evolution events among loci across multiple patients, which has previously been shown in intrahost studies of *Burkholderia pseudomallei* (Price et al., 2013), *P. aeruginosa* (Marvig et al., 2015), and *Burkholderia dolosa* (Lieberman et al., 2011a). We hierarchically considered increasing levels of evidence of convergent adaptation as i) mutations arising intrahost in the same loci, including IGR, among multiple patients, ii) non-synonymous mutations occurring on internal braches of the intrahost phylogeny and/or fixing in a sub-population/clade, and iii) significant tests for gene-wide positive selection. Further, we assessed the occurrence of intrahost IGR switching events for their potential role in host adaptation.

We found that mutations had arose among three intrahost population in 11 genes, three of which, *odhA, mprF*, and *clpX*, have previously been associated with persistence, and one, transcription-repair coupling factor (mfd) is associated with differences in mutation frequencies among strains (Han et al., 2008). In particular, 2-oxoglutarate dehydrogenase (odhA), which is a component of the tricarboxylic acid (TCA) cycle, has been shown to be up-regulated in biofilms (Resch et al., 2005), and *odhA* mutants have been shown to demonstrate attenuated virulence and increased survival in oxygen-limited conditions (Gaupp et al., 2010). Here, we found that in addition to nucleotide variation in *odhA* in three patients, a non-synonymous mutation (S622G) was fixed in the dominant lineage of Patient 2’s intrahost population; the same lineage had a fixed non-synonymous mutation in *clpX* (A82V), providing evidence for genotypic differences involved in clonal interference in at least one patient. Functionally, ATP-dependent protease ATP-binding subunit ClpX is essential for stress tolerance, replication, and biofilm formation, and *S. aureus* ClpX mutants have attenuated virulence (Chen et al., 2015; Frees et al., 2004). Unfortunately, we are unable to assess the phenotypic affect of these mutations, which have not been previously documented. Last, multiple peptide resistance factor (mprF/fmtC) is a membrane protein that regulates the charge of the cell membrane, and mutations in mprF are associated with daptomycin resistance and persistence phenotypes through increased host evasion and adhesion (Richards et al., 2015; Yang et al., 2009). Mutations occurred in *mprF* among three patients; however, non-synonymous mutations were limited to terminal branches of the tree. Overall, the identification of nucleotide and amino acid variation in these loci, combined with the findings implicating their role in persistence, provides good evidence that there is strong selective pressure *in vivo*.

While identifying loci in which mutations commonly arise intrahost can provide candidates for recent adaptation, assessing evidence of gene-wide diversifying selection on internal phylogenetic branches is a stringent test for convergence. Seven loci were significant in the sample including all intrahost clonal isolates; however, further examination showed that these genes were significantly diverged between ST5 and ST72 taxa and may not be associated with recent adaptation. Comparing ST5 and intrahost samples only, a number of genes had an appreciable proportion of sites with ω values (i.e., the fixation ratio of selected vs. neutral mutants) greater than one. However, only immunoglobulin G binding protein A (SpA), a cell-wall associated protein that functions in host immune evasion (Atkins et al., 2008), was significant among ST5 isolates and Patient 4’s intrahost sample. At least one other study has shown within-host variation of the Protein A in longitudinal samples from CF patients (Kahl et al., 2005). Overall, clear evidence of diversifying selection is scant, and in fact, few genes possessed 0-fold vs. 4-fold Θ values (equivalent to dN/dS) greater than one (Supplemental Figure 5). The majority of tested loci had values less than one (mean=0.32), consistent with previous findings from multiple organisms including *S. aureus* (Golubchik et al., 2013; Lieberman et al., 2011a). While purifying selection is likely relaxed within a host on shorter timescales (Rocha et al., 2006), the scarcity of non-synonymous substitutions found here suggests that chronic *S. aureus* carriage in CF patients is sufficient to purge deleterious mutations. There is likely also tremendous selective pressure on strongly deleterious mutations that would impact the function of proteins associated with the aspects of persistence detailed above. Therefore, the amino acid substitutions we find here that have become fixed in dominant intrahost sub-populations likely confer a phenotypic advantage and should be explored further. Last, as three of the four patients studied here possessed relatively closely related ST5 populations, it is likely that the founding populations were already pre-adapted to persistence. This may have broad implications for MRSA epidemiology as the USA100 strain, the predominant healthcare associated strain of MRSA in the US, is ST5. In future studies, comparing transiently to persistently carried *S. aureus* strains may identify ancestrally acquired traits important for persistence.

Selection has historically been thought to act largely on amino acid changes in coding sequences; however, recently it has been shown that IGRs, which account for ~15% of microbial genomes, may also be acted upon by selection (Thorpe et al., 2017). Further, parallel evolution has also been observed among longitudinal intrahost samples of *P. aeruginosa* from CF infection (Khademi and Jelsbak, 2017). In our sample, we observe at least one instance of the same mutation occurring in the IGR flanking PSMb genes. While these loci are involved in pathogenesis (Wang et al., 2007), we do not attempt to interpret the phenotypic affects, if any, which both occurred on terminal branches on the phylogeny. In addition to nucleotide variation in IGRs, switching events have been shown to act as a mechanism for regulating expression in *S. aureus* (Thorpe et al., 2018). Here, we find evidence of three intrahost switching events, two of which occurred between genes associated with quorum sensing. Without expression data, we are unable to gauge the importance of these events; however, this approach could be used in larger datasets combined with RNA-seq data to investigate the frequency and evolutionary impact of intrahost IGR switches. There is currently limited understanding of IGR variation; however, we find evidence that it may important during persistent *S. aureus* carriage.

Antibiotics exert a strong selective pressure on microbial populations, and CF management often requires prolonged use of broad-spectrum antibiotics based empirically on clinical history and microbiological data (Döring et al., 2012). As antibiotic therapy can vary significantly among patients, it is necessary to assess microbial adaptation in the context of differential resistance profiles among intrahost samples as well as the presence of other important co-colonizing organisms. The patients studied here demonstrated common clinical CF trajectories characterized by the consistent isolation of multiple organisms and subsequent treatment. The ST5 MRSA populations identified in Patients 1, 2, and 4 were nearly all universally resistant to fluoroquinolones, marcrolides, and lincosamides, well-adapted to persisting through frequent antibiotic treatments targeting other organisms. Acquired resistance to gentamicin, trimethoprim/sulfamethoxazole, and tetracycline was observed; yet, despite chronic treatment with inhaled trobramycin and frequent treatment with bactrim, resistance did not sweep in the intrahost populations of Patients 1 and 2. In fact, there were instances in each patient where gentamicin resistant and susceptible isolates were cultured on the same day (Figure 3). Further, the transposon harboring the resistance conferring gene, AAC(6’)-Ie-APH(2’’)-Ia, appeared mobile in both patients, as it was contemporaneously identified in isolates from different clades. This finding has implications for treatment, as the overlap in time of extant resistant and susceptible lineages may complicate the choice of a fully active therapeutic regimen.

The ecology of microbes in the airways of CF patients is complex, posing challenges for treatment and overall management. Chief among those challenges is parsing which cultured organisms pose a clinical threat or may in fact be protective. This requires detailed knowledge of pathogen adaptation during prolonged carriage in the context of treatment and polymicrobial interactions. Even with these data, teasing apart the relative contribution of patient-specific selective pressures – age, antibiotic exposure, pathogen and host genotype, immunity, and microbiome – may be arduous. To further resolve these intrahost dynamics, we carried out a detailed analysis of long-term MRSA carriage. Our findings support previous studies illustrating appreciable intrahost diversity as well as persistence of a dominant lineage. We also show that intrahost adaptation is a continual process, despite purifying selective pressure, and provide targets that should be investigated further for their function in CF adaptation. The role of previously CF adapted *S. aureus* lineages and how those lineages, once established in the CF airways, resist intruders requires further analysis. The ultimate goal is to identify key patient and pathogen predictors of long-term CF outcomes that can be used to optimize therapy to eliminate pathogenic bacteria while protecting the resident protective flora in an effort to improve quality of life.

## Acknowledgements

The authors would like to thank Xavier Didelot for technical assistance with BactDating and Sergei L Kosakovsky Pond for technical assistance with HyPhy. It is a noble deed when creators of analytical tools take the time to assist end-users with analysis questions. The authors would also like to thank Brian J Arnold for his guidance, distilled from hours of stimulating discussions about bacterial population genomics. The work of this paper was supported in part by NIH NIAID grant K23 AI095361 (MZD).

## Author Contributions

TA performed the analysis, constructed figures, drafted the initial manuscript, and revised the final draft. JPR abstracted data from the patient medical records and approved the final draft. ZY performed laboratory analysis and microbiological processing of the isolates and approved the final draft. MZD conceived the study, provided funding for whole genome sequencing, assisted in drafting the initial manuscript, and revised the final draft.

## Data Availability

Whole-genome sequencing data have been deposited in NCBI Sequence Read Archive (SRA) under BioProject PRJNA527261.

