## Supplemental figures and tables for "Long-term intrahost evolution of methicillin resistant *Staphylococcus aureus* among cystic fibrosis patients with respiratory carriage"

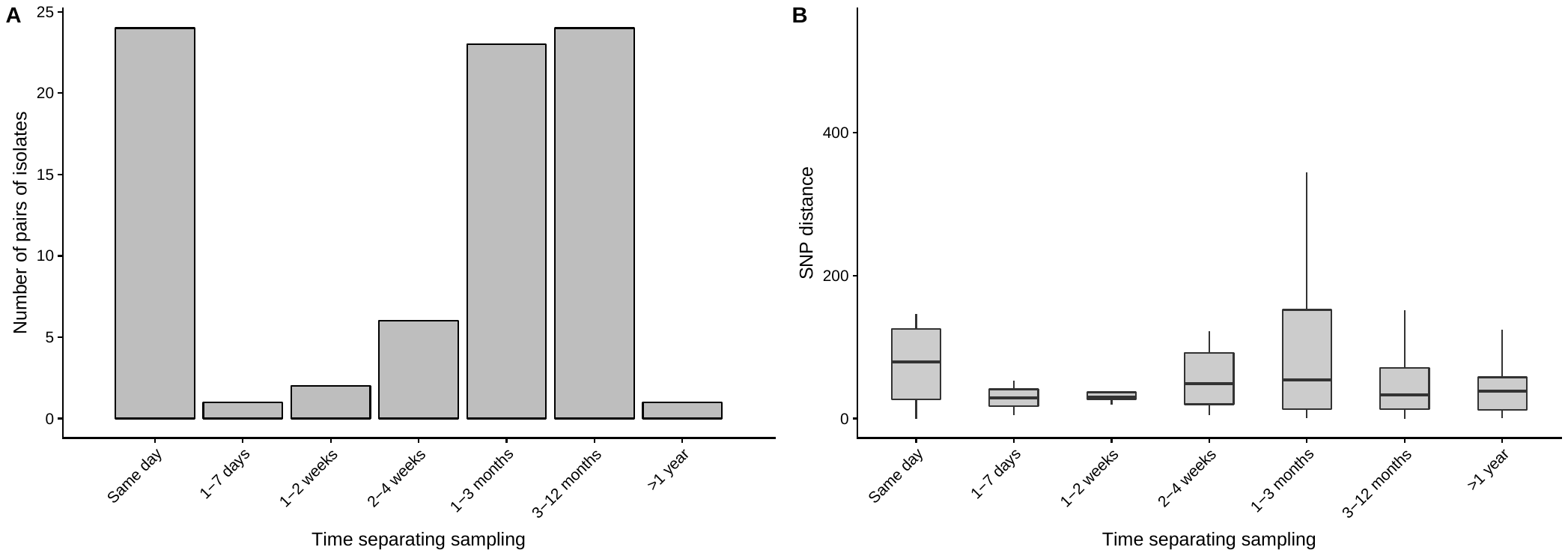

**Supplemental Figure 2. (A) Distribution of days separating pairs of sampled isolates stratified into seven categories; (B) Pairwise SNP distances stratified by categorized duration of time separating collection date of isolates.** As expected, pairwise SNP distances generally increased with time separating the collection date of isolates. This is consistent with what is observed when assessing root-to-tip phylogenetic distance to determine temporal signal. Note the significant range in intrahost diversity observed between pairs of isolates collected on the same day.

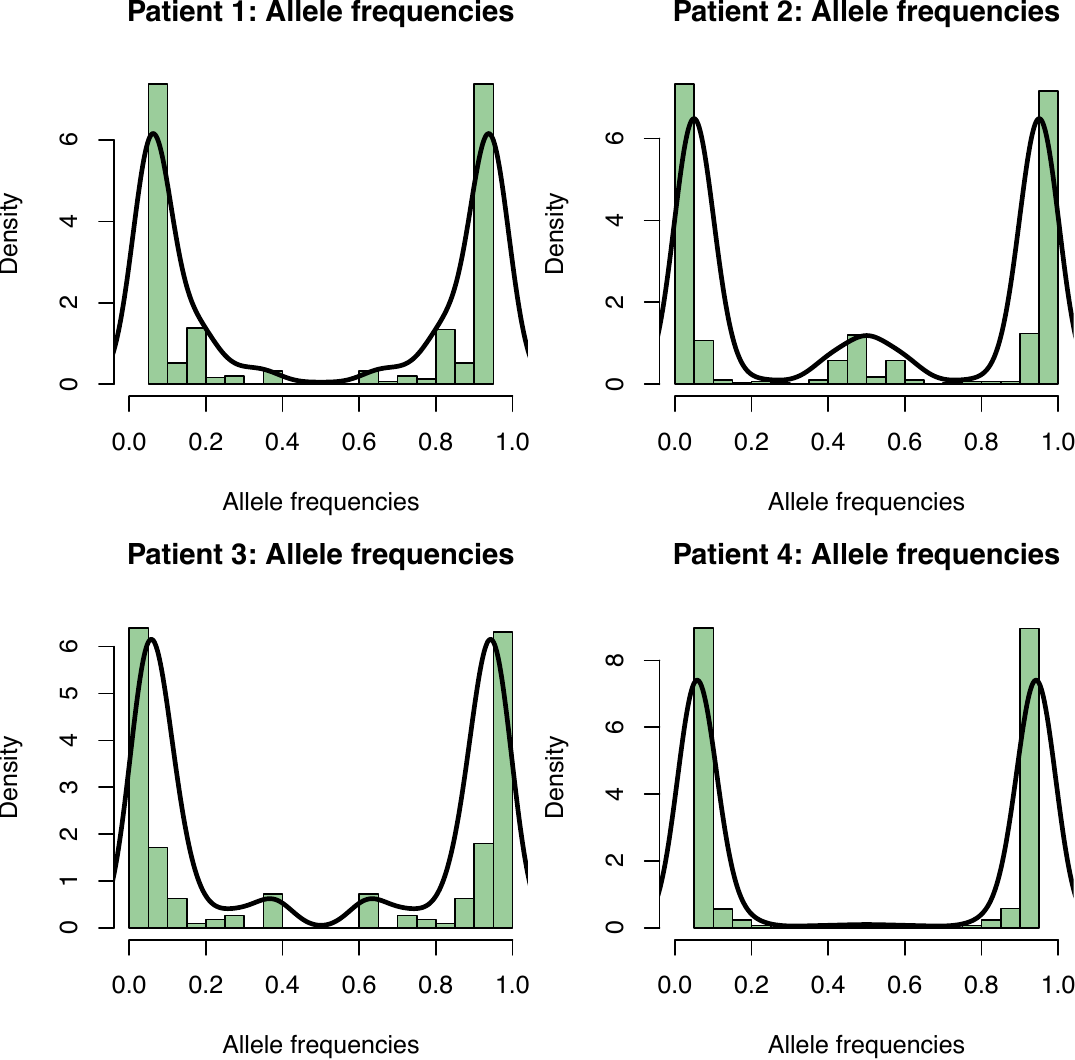

**Supplemental Figure 1. Distribution of allele frequencies among clonal intrahost populations.** The plots are symmetrical and only include the intrahost isolates without reference to illustrate mutations that were intermediate or fixed in the population. For example, Patient 2 has a higher density of intermediate frequency mutations than Patient 4, the effects of which can be seen in the maximum likelihood phylogenies (Figure 2).

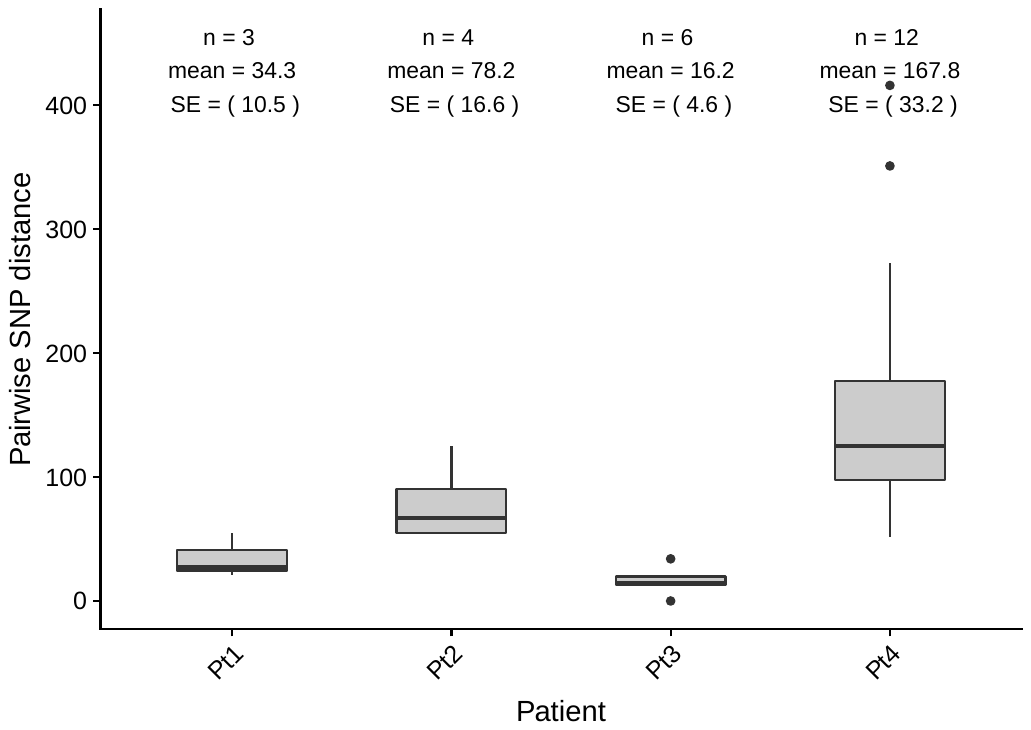

**Supplemental Figure 3. Mean pairwise SNP differences among isolates collected on the same day grouped by patient.** The number same-day isolate pairs, mean, estimates, and standard errors (SE) are given for each patient. Overall, Patient 4 (Pt4) had significantly higher diversity among same-day pairs then other patients.

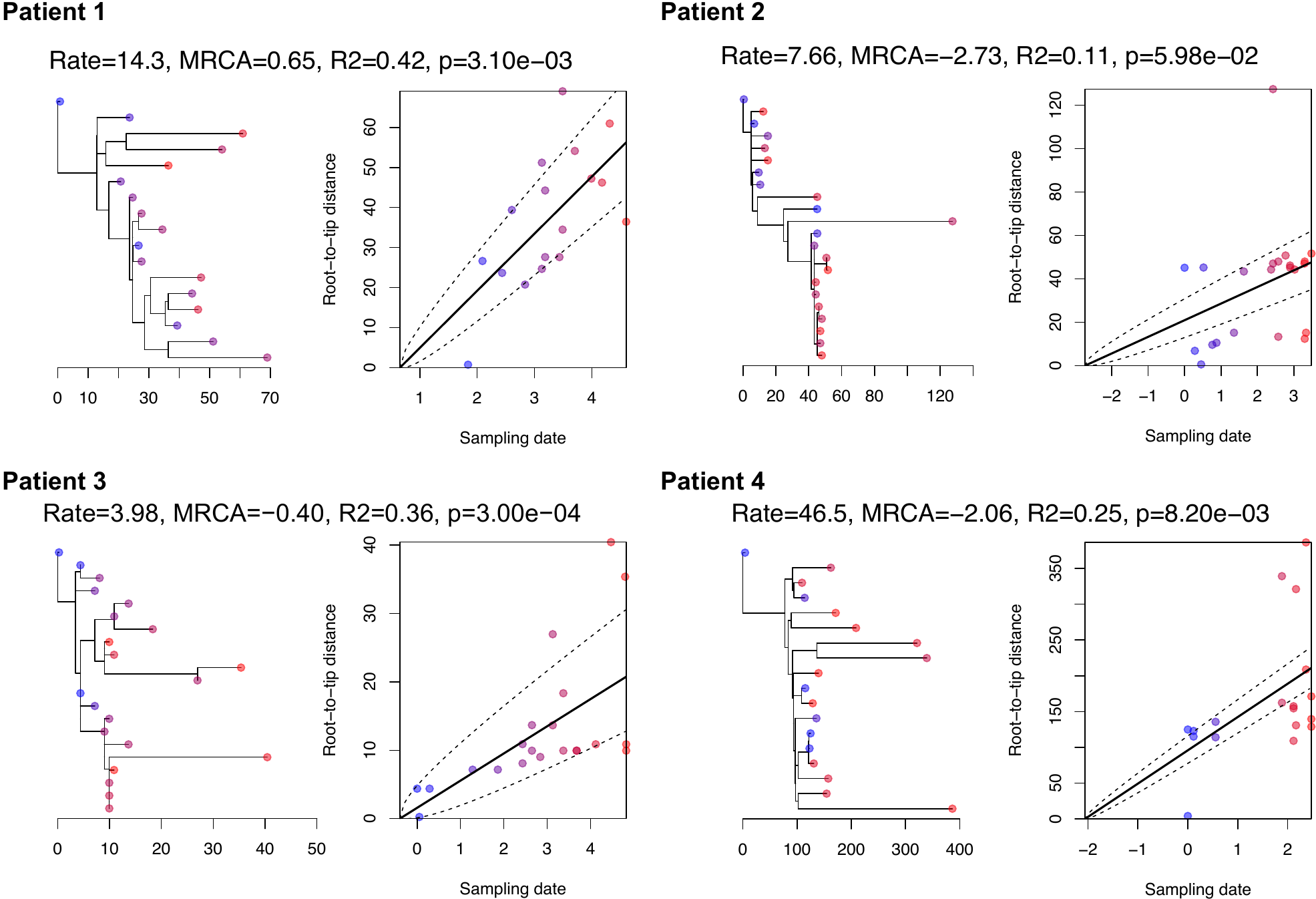

**Supplemental Figure 4. Maximum likelihood core genome phylogenies (left) with best-fit root based on root-to-tip correlation (right) of collection date and genetic distance.** Raw mutation rate in SNPs/genome/year (Rate), most recent common ancestor in years (MRCA), correlation coefficient (R2), and significance test (p) are given at the top of each figure. Isolates are dated based on the day of collection relative to the collection date of the first isolate (day 0). Therefore, clonal populations may have MRCA dates >0 (e.g., Patient 1).

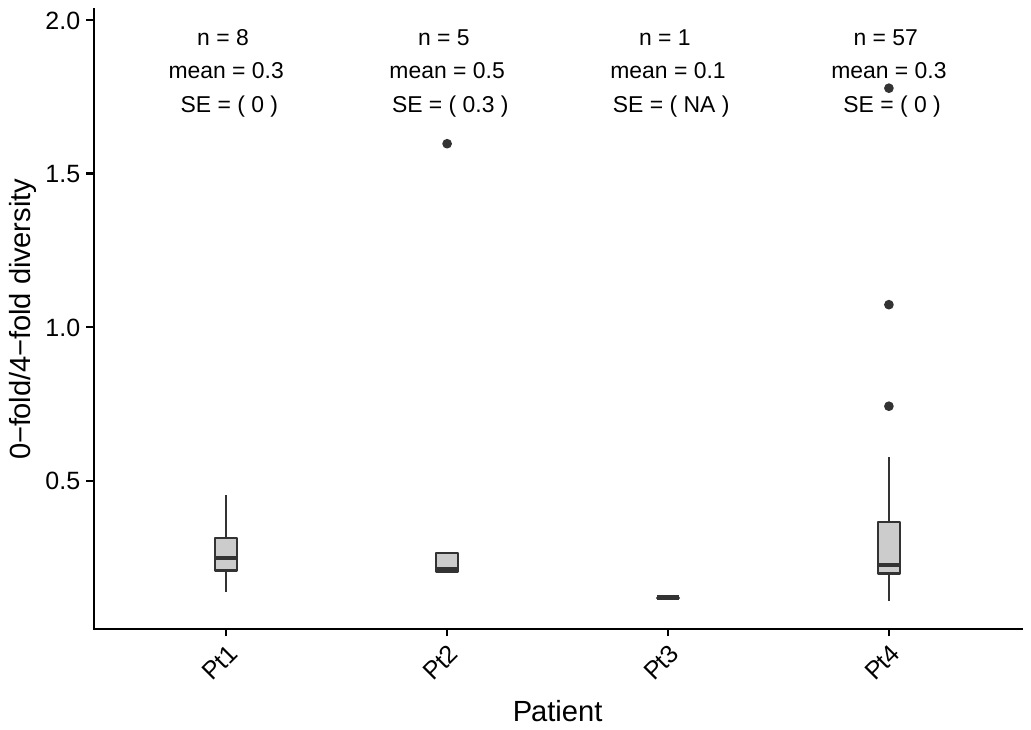

**Supplemental Figure 5. Diversity (Watterson’s Θ) at 0-fold and 4-fold degenerate sites for each patient.**
